# Orchestrating Spatial Transcriptomics Analysis with Bioconductor

**DOI:** 10.1101/2025.11.20.688607

**Authors:** Helena L. Crowell, Yixing Dong, Ilaria Billato, Peiying Cai, Martin Emons, Samuel Gunz, Boyi Guo, Mengbo Li, Alexandru Mahmoud, Artür Manukyan, Hervé Pagès, Pratibha Panwar, Shreya Rao, Callum J. Sargeant, Lori Shepherd Kern, Marcel Ramos, Jieran Sun, Michael Totty, Vincent J. Carey, Yunshun Chen, Leonardo Collado-Torres, Shila Ghazanfar, Kasper D. Hansen, Keri Martinowich, Kristen R. Maynard, Ellis Patrick, Dario Righelli, Davide Risso, Simone Tiberi, Levi Waldron, Raphael Gottardo, Mark D. Robinson, Stephanie C. Hicks, Lukas M. Weber

## Abstract

Spatial transcriptomics technologies provide spatially-resolved measurements of gene expression through assays that can either target selected genes or capture transcriptome-wide expression profiles. The complexity and variability of these technologies and their associated data necessitate multi-step workflows integrating diverse computational methods and software packages. We provide a freely accessible, open-source, continuously updated and tested online book containing reproducible code examples, datasets, and discussion about data analysis workflows for spatial omics data using Bioconductor in R, including interoperability with Python.

## Main text

Spatial transcriptomics technologies are widely used in biomedical research including cancer biology, neuroscience, immunology, and developmental biology [1, 2, 3, 4, 5]. These technologies enable the quantification of spatially-resolved gene expression within tissue sections, providing powerful information on tissue organization and interactions between cells. A number of protocols and technologies are available, which differ in their spatial resolution, the number of genes that can be detected, and sensitivity and specificity. Spatial transcriptomics technologies may be grouped into sequencing-based, which capture RNA from an untargeted or transcriptome-scale set of transcripts at barcoded spatial locations using a sequencing-based readout, and imaging-based, which use a fluorescent readout to identify individual RNA molecules from a typically targeted set of transcripts at subcellular spatial resolution and can be aggregated to cellular resolution [2, 3, 4, 5]. Technologies for other modalities, including spatial proteomics [6, 7, 8] and spatial multi-omics [9], provide further views of spatially-resolved molecular and histological features within cells and tissues.

Computational analyses of spatial transcriptomics data consist of a complex sequence of analysis steps, including preprocessing, quality control, intermediate processing, and downstream analyses, which are connected into workflows. Numerous methods are available for each analysis step (e.g. see [4] for a review). A crucial task for data analysts is to select appropriate computational methods for each step given the data type and experimental context, and to connect the inputs and outputs of different methods in a modular manner to build a complete workflow. Most methods are implemented as R or Python software packages. Standardized data structures, such as *SpatialExperiment* [10] (R/Bioconductor), *AnnData* [11] and *SpatialData* (Python), and structures in the *Seurat* [13] and *Giotto Suite* [14] frameworks (R), facilitate connections between methods. Extensions provide additional capacity for data types from specific technologies (e.g. [15, 16]), or to convert data structures between R and Python (e.g. [17, 18]). Preprocessing steps are generally platform-specific, depending on the format of the raw data (e.g. read alignment or cell segmentation). After preprocessing, the data are usually summarized as a gene expression count table, aggregated at the level of spatial locations (e.g. spots, beads, or bins) or single cells. Subsequent analysis steps use the gene expression count table together with the spatial information as the starting point, e.g. for quality control, feature selection, dimensionality reduction, clustering, and differential testing. Many of these analysis steps are adapted from single-cell RNA sequencing analysis workflows (e.g. [19]), with adaptations to the properties of spatial data such as taking into account distances between observations and considering the number of cells per spatial location. Various downstream analyses, e.g. spatially-aware cell type compositional and interaction analyses, are also applicable to spatial proteomics and other spatial omics data [7, 8, 9].

Bioconductor is a long-standing community-based project that aims to develop and share open-source R-based software for high-throughput biological data analysis [20, 21, 22] (Figure S1). The Bioconductor project started in 2001, and has grown to include more than 2,400 software packages (as of the April 2026 release; Figures S2-S3). Software packages are contributed by numerous research groups around the world, while the overall project and core infrastructure are coordinated and maintained by the Bioconductor Core Team, advised by Community, Technical, and Scientific Advisory Boards. Bioconductor components are primarily developed as R packages, with extensions facilitating interoperability with Python. Since software packages and infrastructure are developed by various research groups, these can include the latest state-of-the-art methods and tools, thus providing a rich, flexible, and modular analysis framework for end users. Bioconductor packages undergo continuous build testing, which notifies package maintainers of any installation or runtime errors. Notably, users and developers benefit from documentation requirements and code review, community-based forums, and educational resources [23]. Bioconductor-based workflows can also incorporate R packages from the Comprehensive R Archive Network (CRAN), providing access to an extensive history of R packages implementing advanced statistical methods in areas including (generalized) linear modeling and spatial statistics, machine learning tools, and sophisticated graphical visualization tools.

Here, we provide a freely accessible, open-source resource consisting of an online book containing reproducible code examples, datasets, and discussion about analyses of spatial transcriptomics data using Bioconductor. OSTA is designed for computational analysts, bioinformaticians, and biologists who need to understand, build, and run spatial omics workflows using existing software tools. The goal is to teach the practical structure of these workflows: how different technologies shape input data, how analysis steps connect, which Bioconductor and related packages are available, and how reproducible workflows can be built and adapted for biological questions. Curated pointers to complementary scientific literature and practical resources aim to support this further. Book chapters cover individual analysis steps as well as extended workflows, using downloadable datasets from several commercially available technologies. The book is hosted on the Bioconductor website, and the code examples are regularly tested on the Bioconductor build system on several operating systems, ensuring reliability, stability, and long-term accessibility for users. The code examples use R packages from either Bioconductor or CRAN, and some chapters further demonstrate inter-operability with Python packages from PyPI using *reticulate* [24] and/or *basilisk* [25]. Datasets used in the examples are stored remotely and can be downloaded using functions provided in a companion Bioconductor package *OSTA*.*data* [26] (Table S1). Figure 1 provides a schematic overview of the book content, and Figure 2 summarizes an illustrative application example for a human colorectal cancer dataset [27], demonstrating analysis steps in OSTA across workflows for the 10x Genomics Visium and Visium HD platforms. Figure S1 illustrates how OSTA fits within the Bioconductor and wider R and Python analysis ecosystems. Additional details and workflow summaries for application example datasets from several platforms are provided in Figures S4-S8 and Online methods. Table S2 summarizes available workflows, and Table S3 summarizes technologies, datasets, and methods covered in the book; and, methodology and software underlying each chapter are detailed in the Supplementary Note.

**Figure 1:**
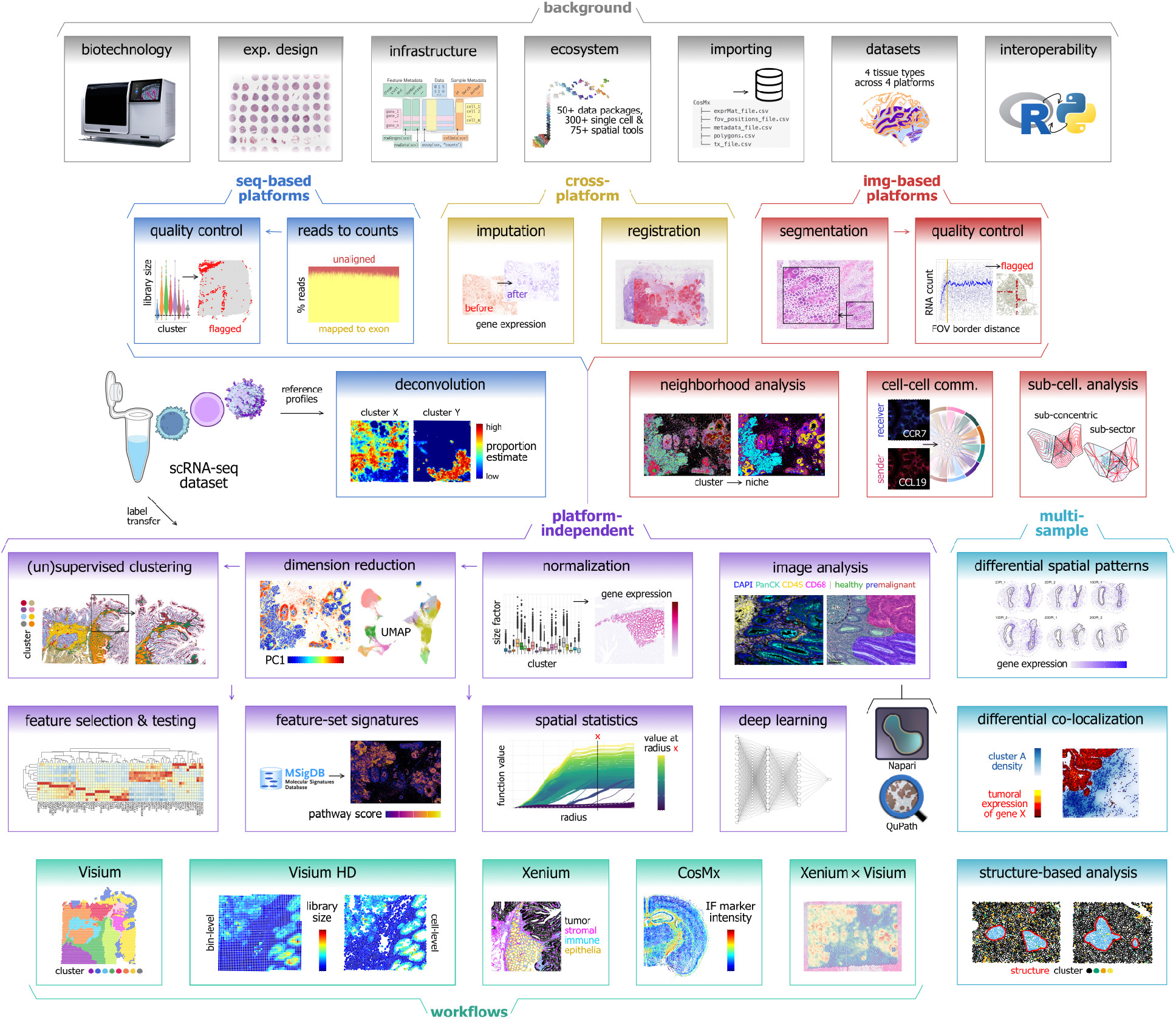
Schematic overview of Orchestrating Spatial Transcriptomics Analysis with Bioconductor (OSTA) book content. The OSTA book consists of a series of chapters grouped into parts, including introduction and background (gray), analyses applicable to sequencing-based technologies (dark blue) and imaging-based technologies (red), platform-independent analyses (purple), multiple-sample analyses (light blue), and cross-platform analyses (yellow). Individual chapters cover individual analysis steps as well as extended workflows for datasets from several major technologies (cyan). Reference single-cell RNA sequencing data may be used for deconvolution of sequencing-based data, and (semi-)supervised clustering of any data; image features may be extracted from, for instance, immunofluorescence or hematoxylin and eosin (H&E) stains using Napari and QuPath, respectively. Arrows indicate an approximate order for a computational data analysis workflow, however, numerous alternative methods are available at each step and may require different processing of data. In summary, OSTA offers the building blocks needed to construct modular data analysis workflows that require careful selection of methods by analysts, depending on the data type, experimental design, and biological question.

**Figure 2:**
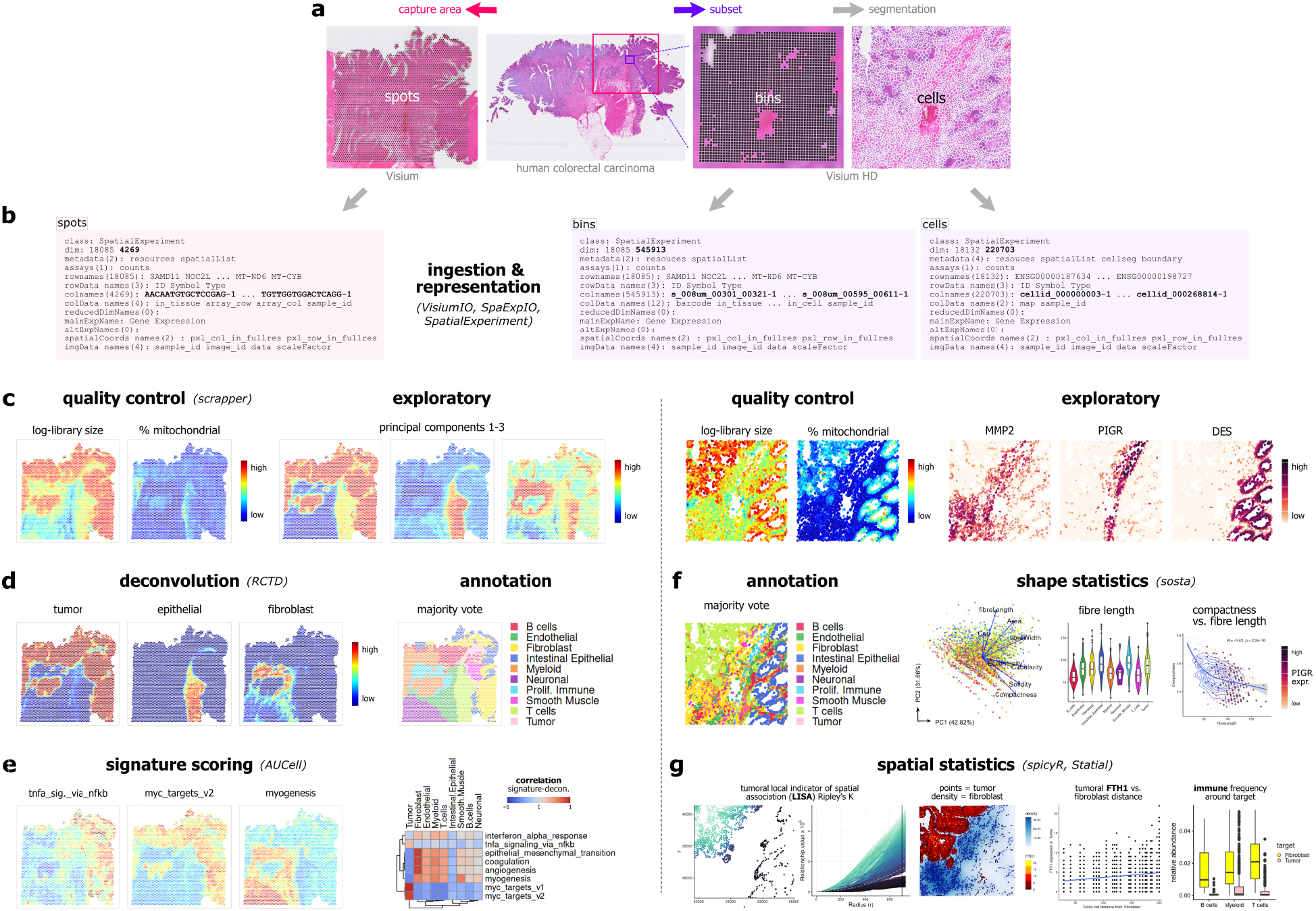
Application example illustrating OSTA analysis workflows applied to Visium and Visium HD data from a human colorectal cancer (CRC) biopsy section [27]. The dataset consists of measurements obtained across multiple spatial transcriptomics technologies from adjacent tissue sections [27]. Here, we demonstrate Visium and Visium HD analyses. Additional details are provided in Figures S4-S7. **(a)** Whole-section hematoxylin and eosin staining contrasting Visium spot-level capture (left) with Visium HD bin-level measurements and cell segmentation (right). **(b)** Observation-level measurements are readily loaded using existing R/Bioconductor tools, and represented as *SpatialExperiment* objects where rows correspond to features and columns to spots, bins, and cells, respectively. **(c)** Platform-specific quality control (QC) and exploratory analyses, including QC metrics, principal components, and spatial marker gene expression. **(d)** Spot-level Visium data are used for cell type deconvolution and annotation of immune, stromal, and tumoral subpopulations. **(e)** Feature-set signature scoring links spatial regions to biological programs and summarizes concordance with deconvolution-derived labels. **(f)** Cell-level Visium HD analyses extend to annotation and morphology-derived shape statistics. **(g)** Spatial statistics quantify local spatial association, proximity-dependent expression, and cell-type neighborhood composition. Together, this example illustrates how OSTA analysis workflows connect platform-specific data types to biologically interpretable spatial analyses across resolutions.

Existing frameworks and tutorials for data analysis workflows for spatial transcriptomics include *Seurat* [13], *Giotto Suite* [14], *Museum of spatial transcriptomics* [4], and *Voyager* [15] (in R), and *Squidpy* [28] (in Python) (Figure S3). Our guiding philosophy is that effective spatial omics analysis requires both conceptual orientation and executable examples; we therefore emphasize modular workflows, common data structures, reproducible code, public datasets, and practical examples, rather than providing formal mathematical foundations. A key advantage of our approach is that both the overall resource and the included methods and tools are developed by various research groups from multiple institutions and countries, thus ensuring that we have included a wide range of representative state-of-the-art scientific methods and analysis approaches. We also emphasize R-Python interoperability with examples in several chapters. In addition, the modularity of the Bioconductor ecosystem allows users to easily adapt our workflows to include new methods, and the continuous Bioconductor build testing ensures that examples remain error-free, while the Bioconductor support site and community forums provide accessible venues for users to ask questions. The development version of the book is hosted on GitHub, which enables additional continuous testing using a GitHub Actions workflow, and provides an additional interface for users to submit issues, provide suggestions and feedback, and contribute content.

Other existing resources provide code examples, tutorials, and discussion on guidelines for analyses of single-cell RNA sequencing data, with extensions to spatially-resolved data, including *Orchestrating Single-Cell Analysis with Bioconductor* (OSCA) [19] in R using Bioconductor, and *Single-Cell Best Practices* [29] in Python using *scverse* [30]. By contrast, our book focuses on spatially-resolved omics data, beginning with introductory discussion on data types and using example datasets from several technologies. This allows us to focus in more detail on the methodological issues and available methods for spatially-resolved data. For some analyses, single-cell methods can be repurposed to provide a computationally efficient baseline for method comparisons, which we discuss in the relevant sections. To facilitate long-term accessibility and maintenance, we restrict code examples to methods available as R packages from Bioconductor or CRAN, or Python packages from PyPI. While we also discuss several key methods available from other sources (e.g. packages from GitHub or other non-package code repositories), these are not included within the reproducible code examples. We also do not provide a complete listing of all available methods for each analysis step, instead focusing on widely used methods and those that we have found to be well documented, accessible, and high-performing. Accordingly, OSTA is primarily intended as a user-facing resource for applied analysis with existing tools. It will also be useful for method developers to understand common workflow contexts, data structures, and interoperability requirements, but it is not intended to serve as a textbook on the mathematical or statistical foundations of spatial omics methods. Where relevant, we include references to benchmark evaluation papers, reviews, primary method papers, and other selected related resources comparing additional available methods and providing deeper methodological details. Our resource is intended as a community-driven, living document that will be updated and extended to cover new methods, data types, and technologies as they become available. The book is available from Bioconductor at https://bioconductor.org/books/OSTA/, and we welcome suggestions, feedback, and contributions from the spatial omics research community.

## Online methods

### OSTA book infrastructure, hosting, and testing

The *Orchestrating Spatial Transcriptomics Analysis with Bioconductor* (OSTA) book is built using open-source publishing tools including *Quarto, R Markdown, bookdown*, and the *BiocBook* [31] Bioconductor package. The rendered version of the book is hosted on the Bioconductor website at https://bioconductor.org/books/OSTA/, and the source code is publicly accessible from Bioconductor and GitHub. The initial version of the book was released as part of Bioconductor version 3.22. The book infrastructure and overall approach are based on and extend previous related Bioconductor resources including *Orchestrating Single-Cell Analysis with Bioconductor* (OSCA) [19].

Consistent with standard Bioconductor package development guidelines, we maintain separate *release* and development (*devel* ) versions, where the release version is relatively stable and intended for use by most readers, and the development version incorporates latest updates and extensions. The release version is updated to match the development version approximately every six months. Both versions are regularly tested on several operating systems using the Bioconductor Build System infrastructure (up to three times per week), ensuring that all code examples run error-free, and dependency packages are accessible. In addition, we maintain a GitHub Actions workflow in the GitHub repository to run tests when updates are made to the source code. The GitHub page also facilitates interaction with the wider community, allowing users to submit issues to provide suggestions and feedback, as well as code contributions.

### OSTA book structure and content

The OSTA book is structured as a series of parts and chapters containing reproducible code examples and text discussion on analyses for spatial omics data. The chapters include discussion chapters including introductory and background material, analysis chapters that each cover a specific analysis step, and workflow chapters that contain extended workflows for datasets from several major technological platforms. The chapters are organized into parts relating to certain types of technologies (e.g. sequencing-based and imaging-based platforms) or types of analyses (e.g. platform-independent, multiple-sample, and cross-platform analyses). The reproducible code examples make use of downloadable datasets stored in an online repository, which can be accessed programmatically using functions provided in the companion Bioconductor package *OSTA*.*data* [26]. An overview of available datasets is provided in Table S1. This structure reflects the practical goal of the book: to help readers including computational analysts, bioinformaticians, and biologists move from understanding spatial omics data types to building complete, reproducible analysis workflows using robust computational tooling.

This section provides an overview of the parts and chapters in the current version of the OSTA book (as of July 2026). The chapters are organized into the following parts, listed below. The complete content, including recent updates, can be viewed in the online version of the book. For a schematic overview, see Figure 1. Visual summaries for a representative selection of workflows (Visium, binned/segmented Visium HD, Xenium, and CosMx) are additionally provided in Figure 2 and Figures S4-8. Table S2 summarizes workflows, and Table S3 provides a summary of the technologies, datasets, and methods covered in the book; chapter-wise contents are summarized in a Supplementary Note as well. These examples include colorectal cancer workflows for Visium, Visium HD, and Xenium, and a mouse brain workflow for CosMx, each emphasizing analyses that are specific to the corresponding platform’s spatial resolution and molecular coverage.

- *Background* : introductory material on spatial omics, file formats, data representations, infrastructure and analysis frameworks (in R/Bioconductor and Python, commercial solutions), and interoperability with Python, including the following chapters: *Introduction, Spatial omics, Experimental design, Infrastructure, Ecosystem, Importing data, Example datasets*, and *Python interoperability*.
- *Sequencing-based platforms* : analyses and workflows for data from sequencing-based platforms, including the following chapters: *Introduction, Reads to counts, Quality control, Intermediate processing, Deconvolution, Workflow: Visium DLPFC, Workflow: Visium CRC, Workflow: Visium HD (binned)*, and *Workflow: Visium HD (segmented)*.
- *Imaging-based platforms*: analyses and workflows for data from imaging-based platforms, including the following chapters: *Introduction, Segmentation, Quality control, Intermediate processing, Neighborhood analysis, Cell-cell communication, Sub-cellular analysis, Workflow: Xenium*, and *Workflow: CosMx*.
- *Platform-independent analyses* : downstream analyses and workflows that are applicable to data from both types of platforms, including the following chapters: *Normalization, Dimensionality reduction, Clustering & annotation, Feature selection & testing, Feature-set signatures, Spatial statistics, Image analysis*, and *Deep learning* .
- *Multiple-sample analyses*: analyses and workflows applicable to datasets consisting of multiple samples (e.g. multiple tissue sections), including the following chapters: *Differential spatial patterns, Differential colocalization*, and *Structure-based analysis*.
- *Cross-platform analyses* : downstream analyses and workflows to integrate or combine information across platforms, including the following chapters: *Spatial registration, Imputation*, and *Workflow: Xenium × Visium*.
- *Beyond this book* : topics and analysis tasks not covered in the current version of the book, including references for further reading on these topics.
- *Appendices*: guidelines for *Contributing, Citation* details (including chapter-wise references), *Acknowledgments*, and *Session information* (including chapter-wise runtimes).

### Application examples and workflows

Throughout OSTA, we showcase individual tools and extended workflows through several example datasets, spanning three species (mouse, human, axolotl), seven technological platforms (Chromium; Visium, Visium HD, and Stereo-seq; Xenium and CosMx; imaging mass cytometry), and five tissue types (brain; pancreas; breast, colorectal, and lung cancer). These examples are included to make analysis decisions concrete, such as how quality control, normalization, deconvolution, annotation, feature testing, and spatial statistics depend on the technological platform and biological question.

Different datasets serve to demonstrate specific analysis tasks, e.g., chapters on *multi-sample analyses* rely on datasets that include multiple regions of interest, tissue sections, and/or experimental conditions. Datasets with multiple sections or modalities have been acquired through either disjoint, adjacent-, or same-section measurements. Several chapters additionally rely on single-cell reference data (Chromium) for specific analysis tasks, such as label transfer and deconvolution. All datasets are publicly accessible (see Data availability).

The colorectal cancer dataset from [27] is used repeatedly in sequencing- and imaging-based workflows: Visium provides spot-level analyses, Visium HD supports bin- and segmentation-derived cell-level analyses, and Xenium provides targeted cell-level measurements from the same study. This allows readers to compare how analyses including quality control, deconvolution or cell annotation, and downstream analyses differ across platforms owing to differences in spatial resolution and molecular coverage (Figure 2; Figures S4-S7). Other application examples broaden the biological context beyond colorectal cancer. The CosMx workflow uses a mouse brain coronal section to demonstrate analyses specific to these data (Figure S8); the cross-platform workflow uses paired Visium and Xenium human breast cancer data to demonstrate spatial alignment, integration, and imputation across data layers.

Workflow chapters combine selected methods into extended pipelines, which focus on data aspects that distinguish computational analyses for representative technological platforms. For sequencing-based platforms, we distinguish between analysis at supra-, sub-, or cellular resolution. For imaging-based platforms, we distinguish between Xenium and CosMx, reflecting differences in quality control considerations (and in some cases plexity) between these platforms. The cross-platform workflow relies on Visium- and Xenium-based readouts from adjacent tissue sections. Workflow chapters included in the current version of OSTA (as of July 2026) are listed in the preceding section.

### R-Python interoperability

The OSTA book is developed primarily using the R programming language, and the majority of the reproducible code examples are run in R, using R packages available from either Bioconductor or CRAN. However, we also include several chapters demonstrating interoperability with Python, in order to demonstrate how to include methods available as Python packages within primarily R-based workflows. This interoperability focus is intended to help readers build practical workflows that combine maintained tools across ecosystems while preserving reproducibility and accessibility. These chapters use *reticulate* [24] and/or *basilisk* [25] to set up and manage Python environments, and use Python packages available from PyPI for the analyses. The code examples demonstrate how to convert data objects between R (*SpatialExperiment* [10]) and Python (*AnnData* [11]) using the *zellkonverter* [18] package; alternatively, *anndataR* [17] could also be used. These examples are intended to give readers a starting point for building extended workflows that seamlessly integrate both R and Python-based tools.

## Code availability

The OSTA book is freely accessible from the Bioconductor website at https://bioconductor.org/books/OSTA/, and the source code is available from Bioconductor as well as GitHub at https://github.com/lmweber/OSTA. Software packages used for data analyses within the reproducible code examples are available from Bioconductor, CRAN, and PyPI.

## Data availability

All datasets and metadata used within the reproducible code examples and workflows are publicly accessible. Several datasets are available from Bioconductor’s *ExperimentHub* (EH) database as R objects; these can be queried and retrieved using the EH interface. Additional datasets have been deposited as ‘flat files’, in accordance with their commercial distribution by the corresponding vendors, through an Open Science Framework (OSF) repository; these can be downloaded programmatically using functions provided in the companion Bioconductor package *OSTA*.*data* [26]. For some datasets, supplementary metadata (e.g., deconvolution results, cell type annotations) have been included as well. By default, all data are managed on-disk using *BiocFileCache* to prevent re-retrieval.

## Author contributions

The list of authors is organized as follows:

*Co-first authors*: HLC, YD

*Contributing authors (listed alphabetically)*:

IB, PC, ME, SGun, BG, ML, AMah, AMan, HP, PP, SR, CJS, LSK, MR, JS, MT

*Contributing principal investigator (PI) authors (listed alphabetically)*: VJC, YC, LCT, SGha, KDH, KM, KRM, EP, DRig, DRis, ST, LW

*Co-senior authors*:

RG, MDR, SCH, LMW

*Corresponding authors* :

HLC, RG, MDR, SCH, LMW

Author contributions following CRediT taxonomy:

*Conceptualization*:

HLC, LCT, MDR, SCH, LMW

*Data curation*:

HLC, YD, MDR, LMW

*Formal analysis* :

HLC, YD, IB, PC, ME, SGun, BG, ML, AMan, PP, SR, CJS, JS, MT, LCT, DRig, DRis, MDR, LMW

*Funding acquisition*:

HLC, VJC, SGha, KM, DRis, RG, MDR, SCH, LMW

*Investigation*:

HLC, YD, IB, PC, ME, SGun, BG, ML, AMan, PP, SR, CJS, JS, MT, LCT, DRig, DRis, MDR, LMW

*Methodology* :

HLC, RG, MDR, SCH, LMW

*Project administration*: HLC, MDR, SCH, LMW

*Resources*:

VJC, YC, LCT, SGha, KDH, KM, KRM, EP, DRig, DRis, ST, LW, RG, MDR, SCH, LMW

*Software*:

HLC, YD, PC, ME, SGun, BG, ML, AMah, AMan, HP, PP, SR, LSK, MR, JS, MT, VJC, LCT, LMW

*Supervision*:

VJC, YC, LCT, SGha, KDH, KM, KRM, EP, DRig, DRis, ST, LW, RG, MDR, SCH, LMW

*Validation*: HLC, LMW

*Visualization*:

HLC, YD, IB, PC, ME, SGun, BG, ML, PP, SR, CJS, JS, MT, LCT, DRig, DRis, MDR, LMW

*Writing - original draft* : HLC, LMW

*Writing - review & editing* :

HLC, YD, RG, MDR, SCH, LMW

## Acknowledgments

The authors thank Aaron T. L. Lun for feedback, advice, as well as his substantial contributions to the Bioconductor ecosystem; Abby Spangler, Madhavi Tippani, and Brenda Pardo for initial work and discussions on Visium data analysis workflows and preprocessing procedures carried out at the Lieber Institute for Brain Development; Irene Ruano at the National Center for Genomic Analysis, Barcelona, Spain, for contributions on experimental design; and members of the Bioconductor Core Team and the Bioconductor community for computational assistance and advice. In addition, the authors thank readers from the Bioconductor and wider research communities for feedback, suggestions, and contributions.

## Abbreviations

SNSF: Swiss National Science Foundation
CoG: Consolidator Grant
DAF: Donor-Advised Fund
DECRA: Discovery Early Career Researcher Award
NIH: National Institutes of Health
NIMH: National Institute of Mental Health
NIGMS: National Institute of General Medical Sciences
NHGRI: National Human Genome Research Institute
NSF: National Science Foundation

