## Supplementary Material for "Orchestrating Spatial Transcriptomics Analysis with Bioconductor"

[1]National Center for Genomic Analysis, Barcelona, Spain. [2]Biomedical Data Science Center, Lausanne University Hospital, Lausanne, Switzerland. [3]University of Lausanne, Lausanne, Switzerland. [4]Department of Biology, University of Padova, Padova, Italy. [5]Department of Molecular Life Sciences, University of Zurich, Zurich, Switzerland. [6]Swiss Institute of Bioinformatics, Zurich, Switzerland. [7]Division of Biostatistics, Department of Population Health Sciences, University of Utah, Salt Lake City, UT, United States. [8]Bioinformatics and Computational Biology Division, Walter and Eliza Hall Institute of Medical Research, Parkville, VIC, Australia. [9]ACRF Cancer Biology and Stem Cells Division, Walter and Eliza Hall Institute of Medical Research, Parkville, VIC, Australia. [10]Department of Medical Biology, The University of Melbourne, Parkville, VIC, Australia. [11]Channing Division of Network Medicine, Mass General Brigham, Boston, MA, United States. [12]Berlin Institute for Medical Systems Biology, Max-Delbrück-Center for Molecular Medicine in the Helmholtz Association, Berlin, Germany. [13]Fred Hutch Cancer Center, Seattle, WA, United States. [14]School of Mathematics and Statistics, The University of Sydney, Camperdown, NSW, Australia. [15]Sydney Precision Data Science Centre, The University of Sydney, Camperdown, NSW, Australia. [16]Charles Perkins Centre, The University of Sydney, Camperdown, NSW, Australia. [17]Centre for Cancer Research, The Westmead Institute for Medical Research, The University of Sydney, Camperdown, NSW, Australia. [18]Roswell Park Comprehensive Cancer Center, Buffalo, NY, United States. [19]Institute for Implementation Science in Population Health, City University of New York Graduate School of Public Health and Health Policy, New York, NY, United States. [20]Department of Epidemiology and Biostatistics, City University of New York Graduate School of Public Health and Health Policy, New York, NY, United States. [21]Department of Biostatistics, Johns Hopkins Bloomberg School of Public Health, Baltimore, MD, United States. [22]Lieber Institute for Brain Development, Johns Hopkins Medical Campus, Baltimore, MD, United States. [23]Center for Computational Biology, Johns Hopkins University, Baltimore, MD, United States. [24]Department of Genetic Medicine, Johns Hopkins School of Medicine, Baltimore, United States. [25]Department of Biomedical Engineering, Johns Hopkins University, Baltimore, MD, United States. [26]Department of Psychiatry and Behavioral Sciences, Johns Hopkins School of Medicine, Baltimore, MD, United States. [27]Solomon H. Snyder Department of Neuroscience, Johns Hopkins School of Medicine, Baltimore, MD, United States. [28]Johns Hopkins Kavli Neuroscience Discovery Institute, Baltimore, MD, United States. [29]Department of Electrical Engineer and Information Technology, University of Naples "Federico II", Naples, Italy. [30]Department of Statistical Sciences, University of Padova, Padova, Italy. [31]Padua Center for Network Medicine, University of Padova, Padova, Italy. [32]Department of Statistical Sciences, University of Bologna, Bologna, Italy. [33]School of Life Sciences, Ecole Polytechnique Fédérale de Lausanne, Lausanne, Switzerland. [34]Center for Computational Biology, Johns Hopkins University, Baltimore, MD, United States. [35]Malone Center for Engineering in Healthcare, Johns Hopkins University, Baltimore, MD, United States. [36]Department of Biostatistics, Boston University School of Public Health, Boston, MA, United States. * These authors share first authorship. † These authors share senior authorship. *Additional details on author contributions and order are provided under Author Contributions.* ✉ Correspondence: ``, ``, ``, ``, ``

July 24, 2026

**References** **21**

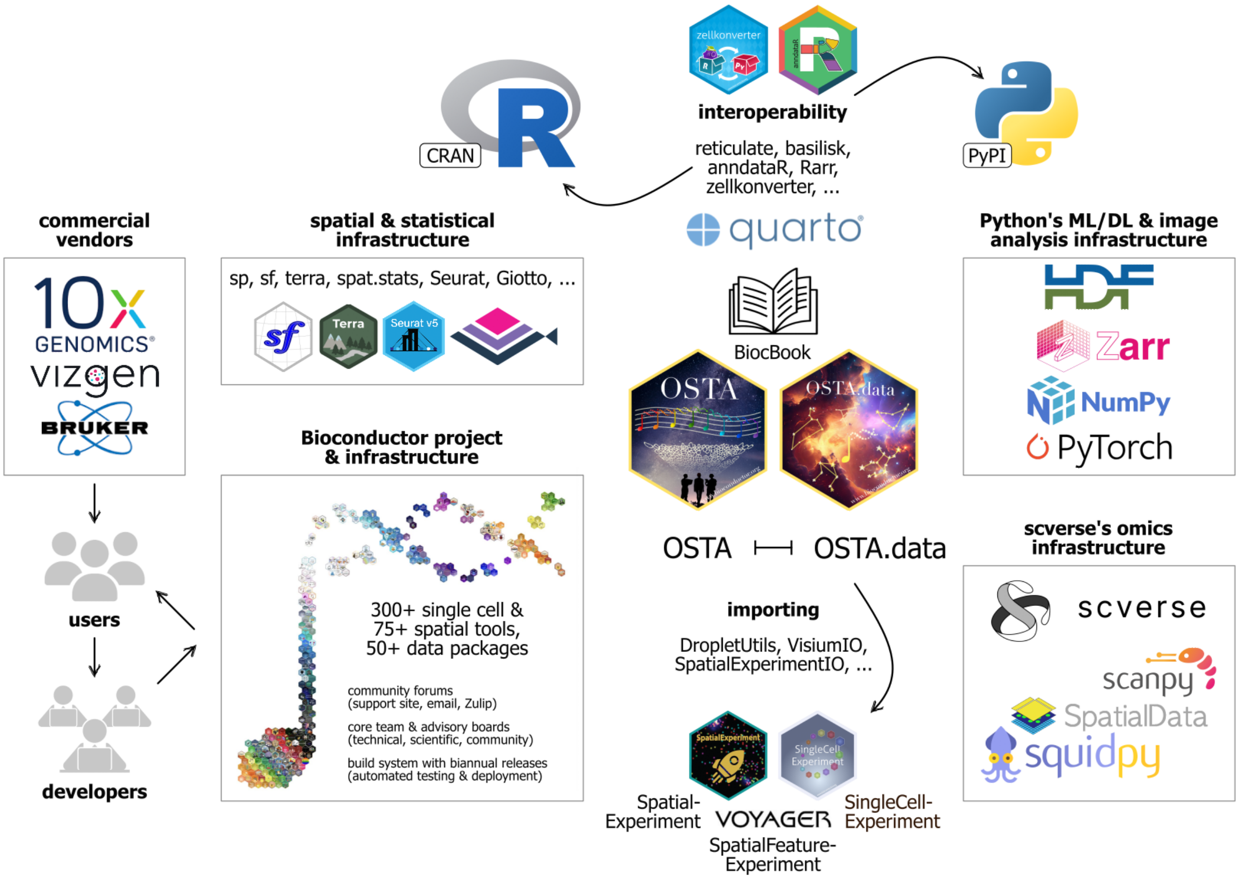

**Figure S1: Schematic illustrating how OSTA fits within the Bioconductor and wider analysis frameworks and ecosystems in R and Python for spatial transcriptomics data.** Data analysis ecosystems comprise tools from many developers and aim to be interoperable, extensible, and adaptable as biological data and computational methods evolve, in addition to offering higher-level supporting infrastructure. Bioconductor offers a suite of software and data packages for single-cell and spatial omics data analysis; project-wide hallmarks include community forums, the Bioconductor Core Team and advisory boards, and an automated build testing system. OSTA relies on various tools for importing and representing data, for rendering and deployment, as well as software that enables interoperability with Python (e.g. data object conversion and running Python code). R-based frameworks, including additional standalone solutions such as *Seurat* [1] and *Giotto Suite* [2], provide access to extensive R packages from the Comprehensive R Archive Network (CRAN) implementing advanced statistical methods (e.g. spatial statistics and linear modeling) and graphical visualization tools, while Python offers rich infrastructure for, in particular, image analysis and machine learning-based applications, as well as frameworks native to the *scverse* ecosystem such as *Squidpy* [3]. In general, technological vendors act as data generators, while users receive data and aim to output research; users may also become developers who, in turn, contribute to the data analysis ecosystems that supply users with the tools and support needed to analyze their data.

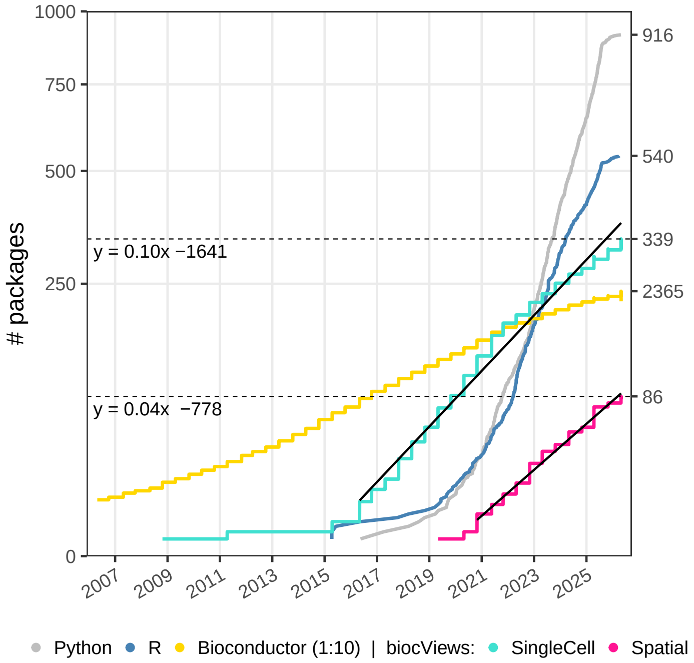

**Figure S2: Dynamics of spatial transcriptomics analysis methods in R and Python, and Bioconductor software.** Cumulative number of methods published for spatial transcriptomics data analysis, in R (blue) and Python (gray), sourced from [4]; and number of Bioconductor software packages, stratified by *biocViews* categories [5] specified by their authors, available for *SingleCell* (cyan) and *Spatial* (pink) analyses, and all categories (yellow; scaled by a factor of 0.1 for clearer visualization). Right-hand side tick marks indicate the number of methods/packages available at the snapshot date, which corresponds to the Bioconductor version 3.23 release date. Black lines and formulas indicate linear model fits for *SingleCell* and *Spatial biocViews*, with slopes corresponding to the average number of packages added per day. Dashed lines indicate the current number of packages available for *SingleCell* and *Spatial biocViews*, as of Bioconductor release version 3.23. Approximately 36.5 *SingleCell* and 14.6 *Spatial* packages are added to Bioconductor each year (slope $\times$ 365.25), with a delay in growth onset of around 4.5 years (difference in x-axis intercepts). The number of R/Python methods corresponds to publications, and Bioconductor statistics were retrieved using *BiocPkgTools* [6], representing a live indication of usable software. [*Data points are 6-monthly; y-scale is square root transformed; snapshot date: April 30, 2026.*]

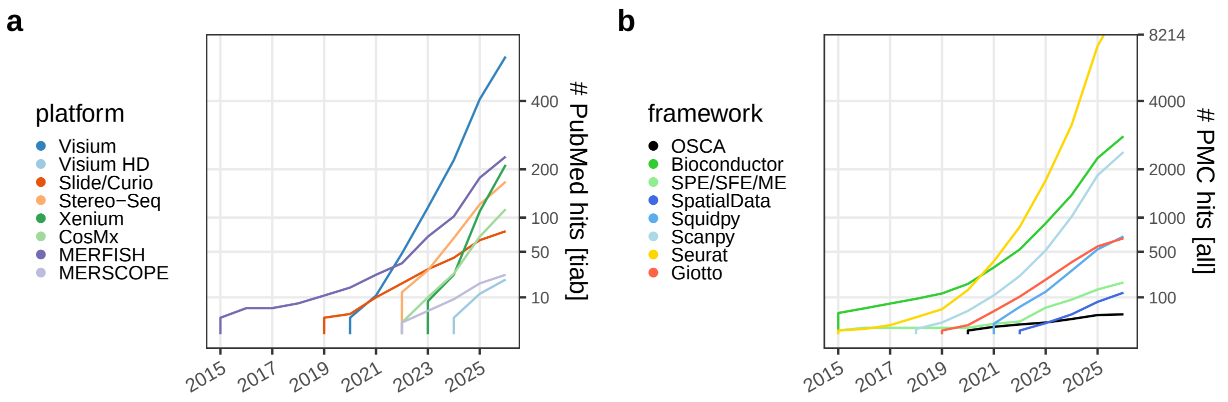

**Figure S3: Dynamics of different spatial transcriptomics technological platform and analysis framework mentions in publications. (a)** Cumulative number of PubMed publication entries that mention specific technological platforms, in addition to "RNA" or "spatial" or "spatially-resolved" or "transcriptomics", in their title or abstract (*tiab*). Slide/Curio includes Slide-seq, Slide-seqV2, and Curio Seeker. **(b)** Cumulative number of PubMed Central (PMC) publication entries that mention specific analysis frameworks, in addition to "spatial transcriptomics" or "spatially-resolved transcriptomics", in their full text (*all*). SPE/SFE/ME includes *SpatialExperiment*, *SpatialFeatureExperiment*/*Voyager*, and *MoleculeExperiment*. OSCA = *Orchestrating Single-Cell Analysis with Bioconductor*. [*Data points are yearly; y-scale is square root transformed; searches were carried out case-insensitive; snapshot date: July 24, 2026.*]

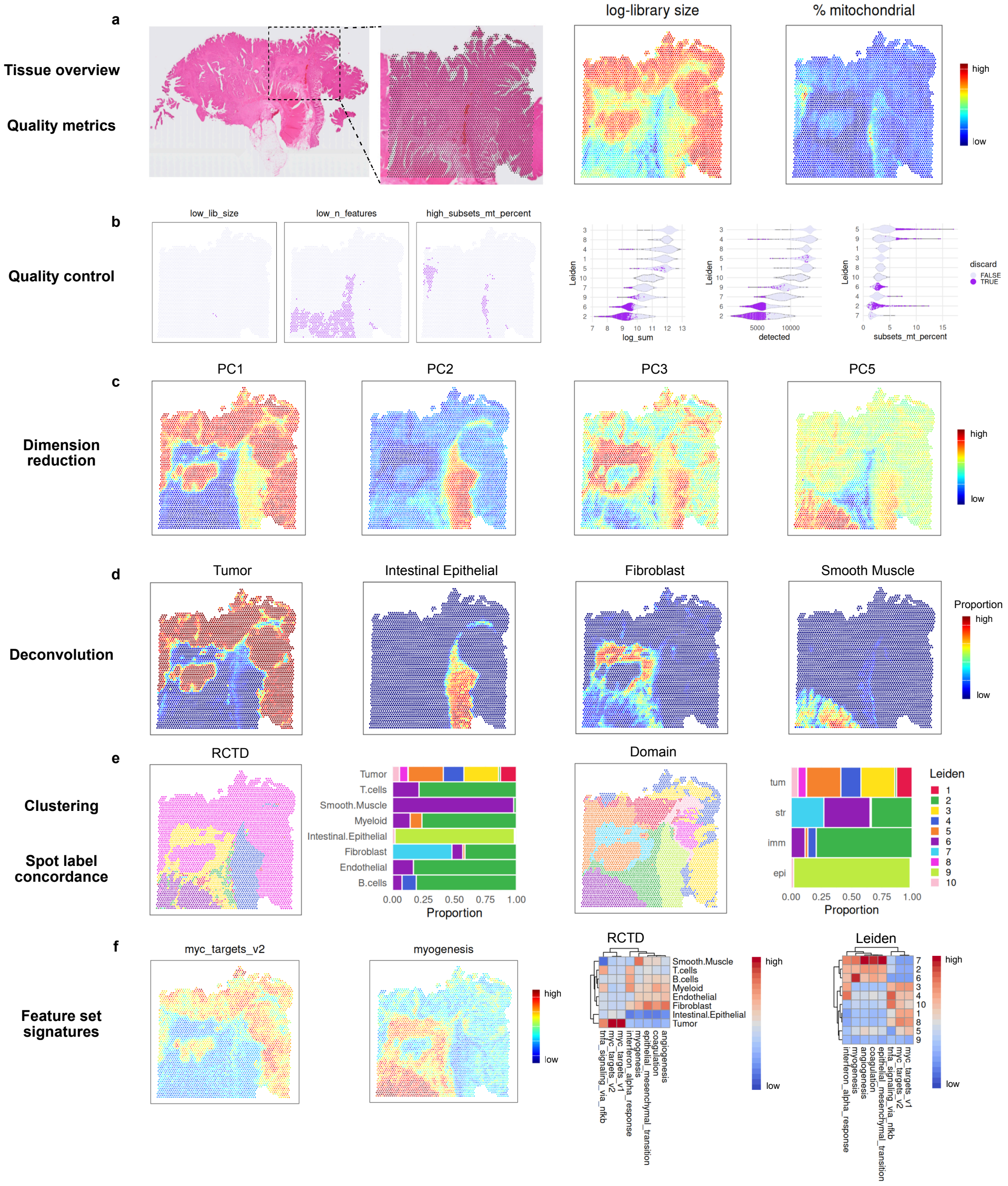

**Figure S4: Visium workflow** using spot-level data on a colorectal cancer (CRC) biopsy section of patient 2 (P2) from [7]. **(a)** Data coverage relative to the tissue-wide H&E staining, and exemplary spot-level quality control metrics. **(b)** Spatial and violin plots highlighting spots flagged as outliers for different metrics. **(c-d)** Selected principal components (PCs) and corresponding deconvolution estimates visualized spatially. **(e)** Comparison of spot label frequencies between deconvolution and clustering results. **(f)** Spatial plots colored by selected feature set signature scores, as well as heatmaps of scores averaged by labels from deconvolution and clustering, respectively.

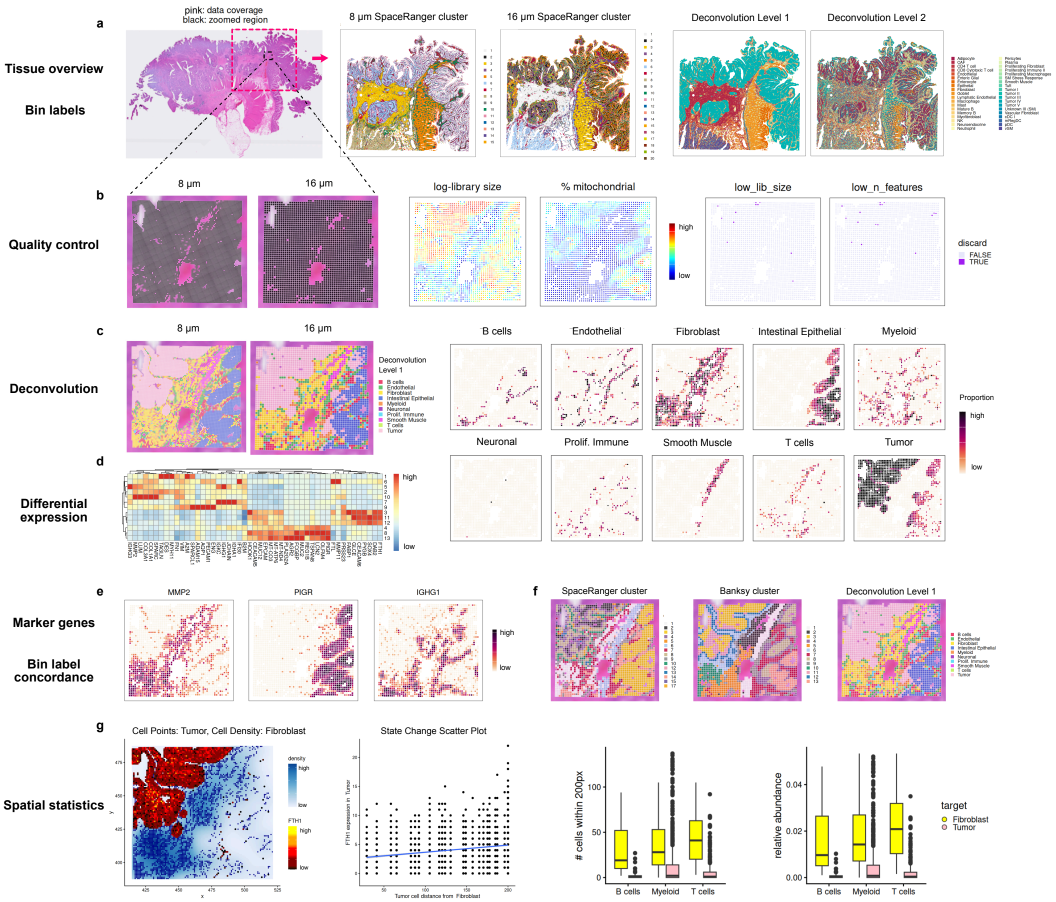

**Figure S5: Visium HD workflow (binned)** using bin-level data on a colorectal cancer (CRC) biopsy section of patient 2 (P2) from [7]. **(a)** Data coverage relative to the tissue-wide H&E staining, as well as clustering and deconvolution labels from SpaceRanger provided by the authors. **(b)** Zoom-in of a region of interest and bin-level quality control using spatially aware metrics. **(c-e)** Clustering and deconvolution at 8 and 16 $\mu$m resolution, respectively, differential expression analysis, and spatial expression of exemplary marker genes. **(f)** Concordance between deconvolution and clustering results. **(g)** Spatial expression of *FTH1* by tumor cells in the context of their proximity to fibroblasts.

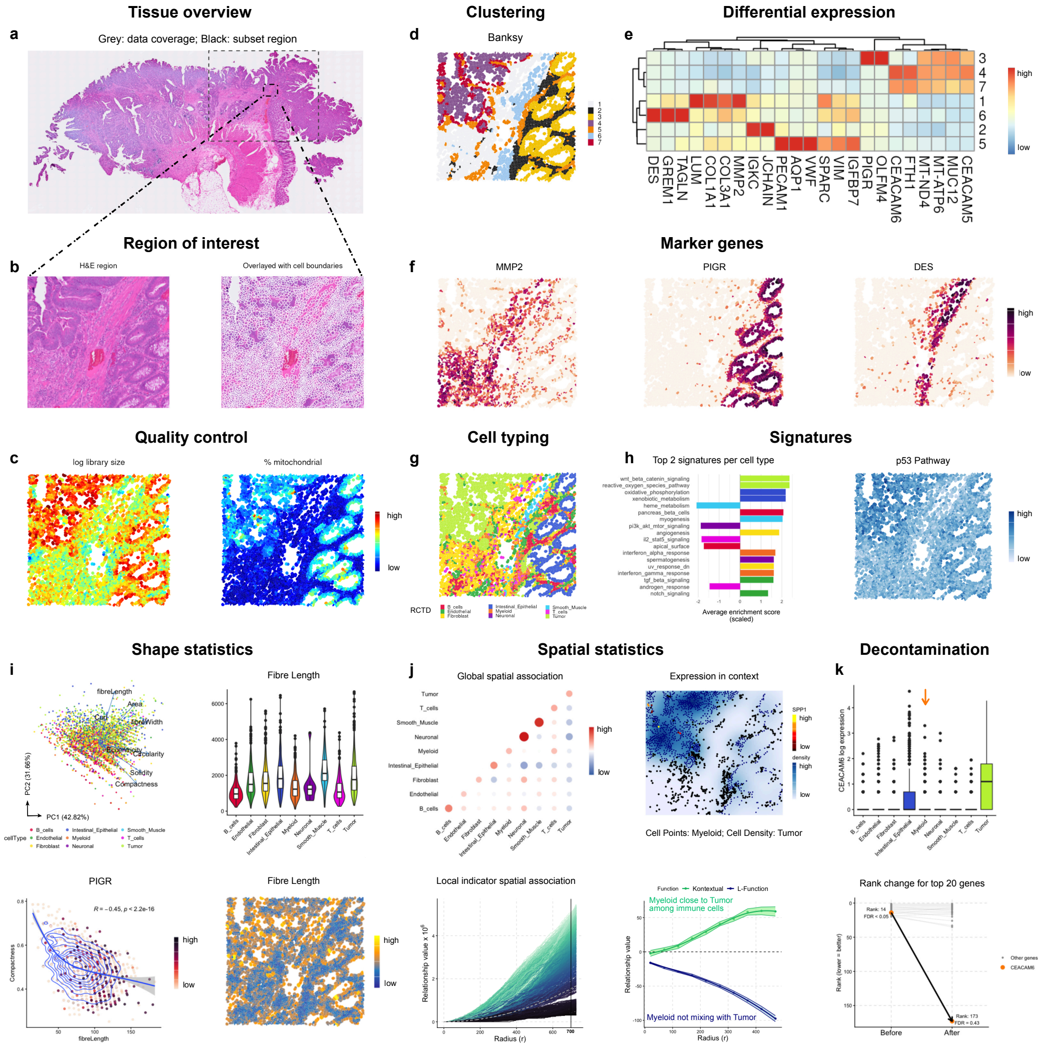

**Figure S6: Visium HD workflow (segmented)** using cell-level data on a colorectal cancer (CRC) biopsy section of patient 2 (P2) from [7]. **(a)** Data coverage relative to the tissue-wide H&E staining. **(b)** Zoom-in of a region of interest. **(c)** Spatial plots colored by exemplary cell-level quality control metrics. **(d)** Unsupervised clustering assignments visualized spatially. **(e)** Heatmap of differential expression analysis-derived marker genes across clusters. **(f)** Expression of selected marker genes for clusters 1, 3 and 6, respectively, visualized spatially. **(g)** Spatial plot colored by assignments from label transfer using matched single-cell reference data. **(h)** Top 2 feature-set signatures per cell type, based on their z-scaled average score; and, p53 pathway scores visualized spatially. **(i)** Principal component analysis on shape metrics and their loadings (arrows); correlation between fibre length and compactness, colored by *PIGR* expression; and, fibre length distribution across cell types and visualized spatially. **(j)** Dotplot of pair-wise cell type co-localization; spatial expression of *SPP1* by myeloid cells in the context of their proximity to tumor cells; *Kontextual* and classical *L*-function capturing the relationship between myeloid and tumor cells for increasing radius. **(k)** Top 20 genes identified as changing in myeloid cells with proximity to tumor cells, before and after decontamination using deconvolution estimates.

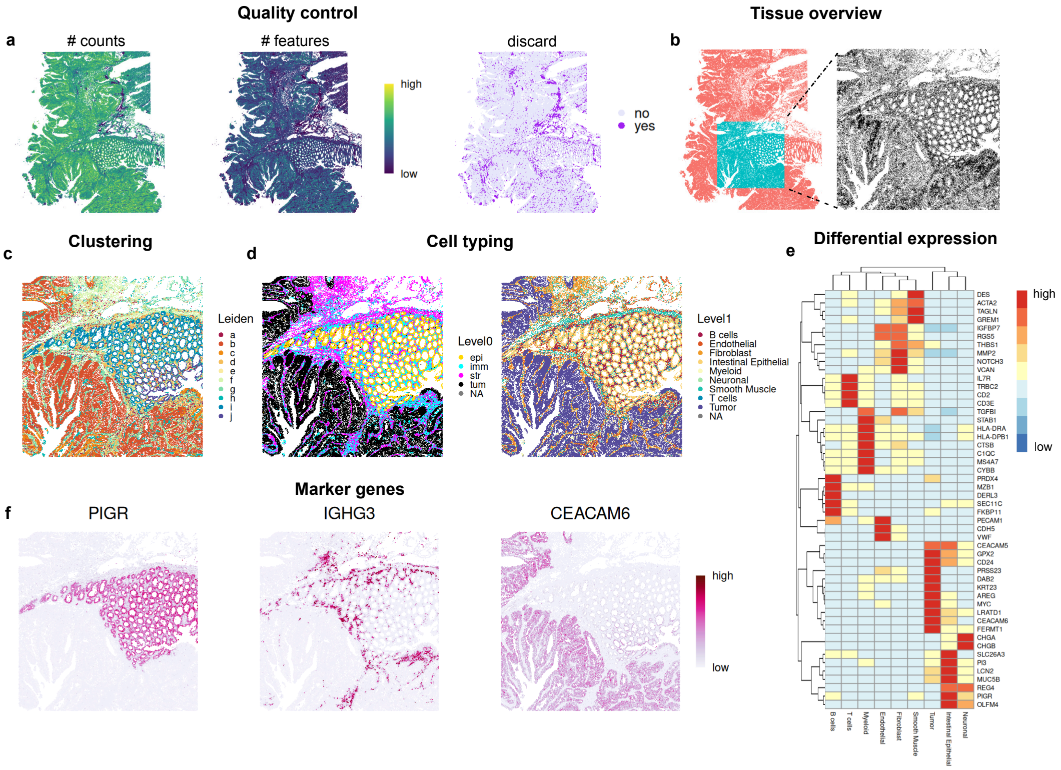

**Figure S7: Xenium workflow** using 422-plex cell-level data on a colorectal cancer (CRC) biopsy section of patient 2 (P2) from [7]. **(a)** Cell-level quality control metrics visualized spatially. **(b)** Zoom-in of a region of interest (to reduce computational burden). **(c-d)** Spatial plots of graph-based clustering assignments (left), label transfer using matched single-cell reference data (right), and annotation into broad biological compartments (middle). **(e)** Heatmap of differential expression analysis-derived marker genes across cluster assignments from label transfer. **(f)** Spatial expression of exemplary marker genes delineating healthy epithelium, IgG+ plasma cells, and cancerous tissue.

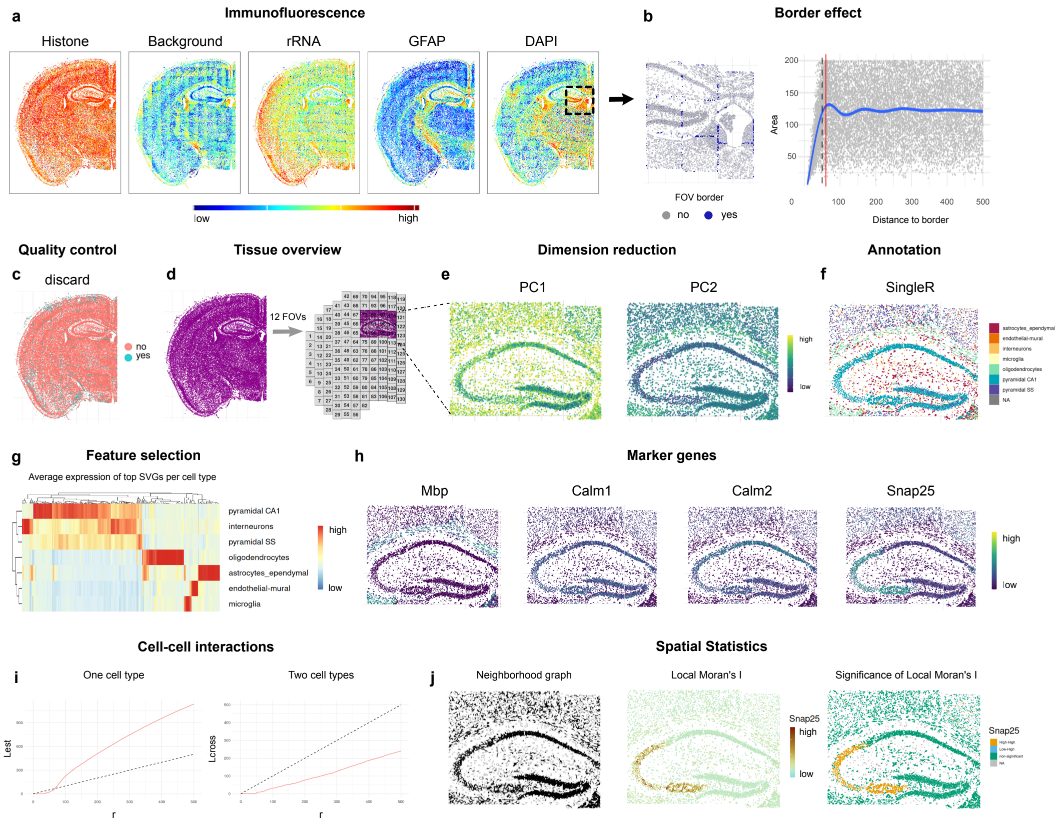

**Figure S8: CosMx workflow** using 1,000-plex cell-level data (Mouse Neuroscience RNA panel) on a mouse brain tissue section (coronal hemisphere) from Bruker. **(a)** Immunofluorescence stainings visualized spatially. **(b)** Zoom-in of a region of interest, highlighting cells flagged as lying on field of view (FOV) borders; and, cell area as a function of FOV border distance. **(c)** Spatial plot highlighting low-quality cells. **(d)** FOV map highlighting a region of interest for subsequent analysis. **(e)** First two principal components (PCs) visualized spatially. **(f)** Cell type annotation with *SingleR* using matched single-cell reference data. **(g)** Heatmap of mean expression values for selected spatially variable genes (SVGs) across cell types. **(h)** Marker gene expression visualized spatially. **(i)** Point pattern analysis of one cell type (pyramidal CA1) and between two cell types (pyramidal CA1 to oligodendrocytes). **(j)** Local Moran's $I$ and its significance level visualized spatially, calculated from a cell-cell neighborhood graph (left-most panel).

| Identifier | Platform | Features | Obs. | Source |
|---|---|---|---|---|
| 1,000-plex CosMx data on **mouse brain (coronal)** FFPE tissue: hemisphere, hippocampus and cortex (Mouse Neuroscience RNA panel) | | | | |
| *CosMx1k_MouseBrain1* | CosMx | 950 | 48,556 | Bruker |
| *CosMx1k_MouseBrain2* | CosMx | 950 | 38,996 | |
| 6,000-plex CosMx data on **human frontal cortex** FFPE tissue (Human 6K Discovery RNA panel) | | | | |
| *CosMx6k_HumanBrain* | CosMx | 6,278 | 188,686 | Bruker |
| Chromium, Visium and Xenium data on consecutive slices of **human breast cancer** | | | | |
| *Chromium_HumanBreast_Janesick* | Chromium | 18,082 | 30,365 | GSM7782698 |
| *Visium_HumanBreast_Janesick* | Visium | 18,085 | 4,992 | GSM7782699 |
| *Xenium_HumanBreast1_Janesick* | Xenium | 313 | 167,780 | GSM7780153 |
| Chromium, Visium (HD) and Xenium data on consecutive slices of **human colorectal cancer** | | | | |
| *Chromium_HumanColon_Oliveira* | Chromium | 18,082 | 279,609 | 10x Genomics |
| *Visium_HumanColon_Oliveira* | Visium | 18,085 | 4,269 | |
| *VisiumHD_HumanColon_Oliveira* | Visium HD | 18,085 | 8,731,400 | |
| *Xenium_HumanColon_Oliveira* | Xenium | 422 | 340,837 | |

**Table S1: Summary of datasets provided through the *OSTA.data* package.** Included are a brief description (species, tissue type), identifier (in R), commercial platform, number of features and observations (obs.), and original source. All data can be queried and retrieved programmatically in R using the *OSTA.data_list()* and *_load()* function, respectively. Datasets have been deposited in an Open Science Framework (OSF) repository at `https://osf.io/5n4q3`; and, data files downloaded with the package are stored and managed in a temporary directory using *BiocFileCache*.

| Visium DLPFC | Visium CRC | Visium HD |
| --- | --- | --- |
| human brain [8]<br>33,538 × 4,992 (*STExampleData*) | human CRC [7]<br>18,085 × 4,269 (*OSTA.data*) | human CRC [7]<br>18,085 × 545,913 (*OSTA.data*) |
| • quality control (*scater*)<br>• log-library size normalization<br>• HVG selection (*scran*)<br>• PCA-based SNN graph<br>• Walktrap clustering (*scran*)<br>• differential expression analysis<br>• interactive analysis (*spatialLIBD*) | • quality control (*scater*)<br>• log-library size normalization<br>• HVG selection (*scran*)<br>• PCA-based SNN graph<br>• Leiden clustering (*igraph*)<br>• RCTD deconvolution (*spacexr*)<br>• feature-set scoring (*AUCell*) | • quality control (*SpotSweeper*)<br>• log-library size normalization<br>• HVG selection (*scran*)<br>• spatially aware clustering (*Banksy*)<br>• RCTD deconvolution (*spacexr*)<br>• neighborhood analysis (*Statial*) |

| Xenium | CosMx | Xenium × Visium |
| --- | --- | --- |
| human colorectal cancer [7]<br>422 × 340,837 (*OSTA.data*) | mouse brain tissue [Bruker]<br>950 × 48,556 (*OSTA.data*) | human breast cancer [9]<br>313 × 167,780 (*OSTA.data*) |
| • quality control (*scater*)<br>• area-based log-normalization<br>• PCA-based SNN graph<br>• Leiden clustering (*igraph*)<br>• label transfer (*SingleR*)<br>• differential expression analysis | • quality control (*SpaceTrooper*)<br>• area-based log-normalization<br>• scRNA-seq label transfer (*SingleR*)<br>• within-cell type SVGs (*DESpace*)<br>• gene expression auto-correlation<br>• cell-cell interactions (*spdep*)<br>• point-pattern analysis (*spatialFDA*) | • spatial alignment (pre-computed)<br>• cell-to-spot aggregation<br>• integration (*harmony*)<br>• joint UMAP embedding<br>• joint PCA-based SNN graph<br>• joint Leiden clustering (*igraph*) |

**Table S2: Summary of workflows presented in *OSTA* v1.2.2.** Included are a brief description (species, tissue type), identifier (in R), commercial platform, number of features and observations (obs.), and original source. All data can be queried and retrieved programmatically in R using the *OSTA.data_list()* and *_load()* function, respectively. Datasets have been deposited in an Open Science Framework (OSF) repository at `https://osf.io/5n4q3`; and, data files downloaded with the package are stored and managed in a temporary directory using *BiocFileCache*.

| | | |
|---|---|---|
| **Biotechnology** | | |
| ST | | [10] |
| Stereo-seq | STOmics | |
| Slide-seqV2 | | [11] |
| Visium | 10x | |
| Visium HD | 10x | |
| Xenium | 10x | |
| CosMx | Bruker | [12] |
| MERSCOPE | Vizgen | |
| MERFISH | | [13] |
| spatial-CITE-seq | | [14] |
| spatial-ATAC-RNA-seq | | [15] |
| DBiT-seq | | [16] |
| SPOTS | | [17] |
| **Infrastructure** | | |
| *SpatialExperiment* | Bioc | [18] |
| *MoleculeExperiment* | Bioc | [19] |
| *SingleCellExperiment* | Bioc | [20] |
| *SpatialFeatureExperiment* | Bioc | [21] |
| *iSEE* | Bioc | [22] |
| *Seurat* | GitHub | [1] |
| *VoltRon* | GitHub | [23] |
| *spatialGE* | GitHub | [24] |
| *Giotto Suite* | GitHub | [2] |
| **Interoperability** | | |
| *Rarr* | Bioc | [25] |
| *alabaster* | Bioc | [26] |
| *basilisk* | Bioc | [27] |
| *anndataR* | Bioc | [28] |
| *zellkonverter* | Bioc | [29] |
| *reticulate* | CRAN | [30] |
| **Datasets** | | |
| mouse brain (Cos) | Bruker | |
| human brain (Vis) | 10x | [8] |
| human breast cancer (Vis, Xen) | 10x | [9] |
| human CRC (Vis/HD, Xen) | 10x | [7] |
| axolotl brain (Stereo-seq) | STOmics | [31] |
| human type I diabetes (IMC) | | [32] |
| *STexampleData* | Bioc | [18] |
| *spatialLIBD* | Bioc | [33] |
| *OSTA.data* | Bioc | [34] |
| **Importing** | | |
| *VisiumIO* | Bioc | |
| *XeniumIO* | Bioc | |
| *SpatialExperimentIO* | Bioc | |

| | | |
|---|---|---|
| **Preprocessing & quality control** | | |
| *stPipe* | Bioc | [35] |
| *Rsubread* | Bioc | [36] |
| *scater* | Bioc | [37] |
| *scrapper* | Bioc | |
| *SpaNorm* | Bioc | [38] |
| *SpotSweeper* | Bioc | [39] |
| *SpaceTrooper* | Bioc | [40] |
| *SegTraQ* | Github | |
| **Dim. red., clustering & anno.** | | |
| *BANKSY* | Bioc | [41] |
| *BayesSpace* | Bioc | [42] |
| *CellAssign* | Bioc | [43] |
| *SingleR* | Bioc | [44] |
| *scType* | CRAN | [45] |
| *Azimuth* | GitHub | [46] |
| *scANVI* | GitHub | [47] |
| *CellTypist* | GitHub | [48] |
| *cellxgenedp* | Bioc | [49] |
| *scDiagnostics* | Bioc | [50] |
| **Neighborhood analysis** | | |
| *RANN* | CRAN | |
| *scider* | Bioc | [51] |
| *hoodscanR* | Bioc | [52] |
| *imcRtools* | Bioc | [53] |
| *DeepST* | GitHub | [54] |
| *GraphST* | GitHub | [55] |
| *NicheCompass* | GitHub | [56] |
| *Novae* | GitHub | [57] |
| *SpaGCN* | GitHub | [58] |
| *STAGATE* | GitHub | [59] |
| **Cell-cell comm.** | | |
| *mistyR* | Bioc | [60] |
| *CCPlotR* | Bioc | [61] |
| *CellChat* | GitHub | [62] |
| *SpatialDM* | GitHub | [63] |
| *SpaOTsc* | GitHub | [64] |
| **Sub-cellular analysis** | | |
| **Segmentation** | | |
| *Bayor* | GitHub | [65] |
| *Proseg* | GitHub | [66] |
| *CellPose* | GitHub | [67] |
| *SSAM* | GitHub | [68] |
| *SPLIT* | GitHub | [69] |
| *segger* | GitHub | [70] |
| *FastReseg* | GitHub | [71] |
| *cellAdmix* | GitHub | [72] |
| **Deconvolution** | | |
| *spacexr* | Bioc | [73] |
| *CARDspa* | Bioc | [74] |
| *SPOTlight* | Bioc | [75] |
| *SpatialDecon* | Bioc | [76] |

| | | |
|---|---|---|
| **Feature sets, selection & testing** | | |
| *msigdbr* | Bioc | |
| *AUCell* | Bioc | [77] |
| *scran* | Bioc | [78] |
| *nnSVG* | Bioc | [79] |
| *DESpace* | Bioc | [80] |
| *spatialDE* | Bioc | [81] |
| **Spat. stat., multi-sample** | | |
| *sf* | CRAN | [82] |
| *sp* | CRAN | [83] |
| *spat.stat* | CRAN | [84] |
| *pasta* | GitHub | [85] |
| *Voyger* | Bioc | [21] |
| *sosta* | Bioc | [86] |
| *spicyR* | Bioc | [87] |
| *Statial* | Bioc | [88] |
| *spatialFDA* | Bioc | [89] |
| **Image analysis** | | |
| *napari* | GitHub | [90] |
| *QuPath* | GitHub | [91] |
| *Prov-GigaPath* | GitHub | [92] |
| *imageTCGA* | Bioc | |
| **Imputation, registration** | | |
| *rliger* | CRAN | [93] |
| *VoltRon* | GitHub | [23] |
| *SLAT* | GitHub | [94] |
| *PASTE* | GitHub | [95] |
| *CeLEry* | GitHub | [96] |
| *SpaOTsc* | GitHub | [64] |
| *STalign* | GitHub | [97] |
| *Tangram* | GitHub | [98] |
| *novoSpaRc* | GitHub | [99] |
| **Further analyses** | | |
| *PRECAST* | CRAN | [100] |
| *harmony* | Bioc | [101] |
| *CellMixS* | Bioc | [102] |
| *slingshot* | Bioc | [103] |
| *monocle* | Bioc | [104, 105] |
| *SpaceFlow* | GitHub | [106] |
| *stLearn* | GitHub | [107] |
| *spaTrack* | GitHub | [108] |
| *CellPLM* | GitHub | [109] |
| *scGPT* | GitHub | [110] |
| *scFoundation* | GitHub | [111] |
| *Geneformer* | GitHub | [112] |
| *SpatialGlue* | GitHub | [113] |
| *MOFA2* | Bioc | [114] |

**Table S3:** Overview of spatial transcriptomics technologies, datasets, and methods that are either mentioned or used throughout the OSTA book (v1.2.2). Methods (cursive) including software implemented in R and/or Python; if available, source (company name or code repository) and reference are also listed. [Bioc = Bioconductor, 10x = 10x Genomics, Vis = Visium, HD = Visium HD, Cos = CosMx, Xen = Xenium, IMC = imaging mass cytometry, CRC = colorectal carcinoma, dim. red. = dimensionality reduction, anno. = annotation, cell-cell comm. = cell-cell communication, spat. stat. = spatial statistics.]

### Supplementary Note

Contents outlined here align with OSTA v1.2.2; introductory chapters are not included.
*Indicates that a chapter is purely discussion-based (i.e., no code is being evaluated).

#### I Background

***Spatial omics.**** OSTA commences with an overview of spatial transcriptomics (ST) technologies, divided into sequencing-based (sST) and imaging-based (iST) approaches, with focus on commercial platforms. Introductory chapters preceding these parts provide additional detail on sST and iST, respectively. Other types of spatial omics (e.g., proteomics, metabolomics, lipidomics) and multimodal approaches are also discussed, as well as efforts towards spatial molecular profiling beyond 2D sections. In particular, reconstruction of "virtual tissue blocks" from serial sections (2.5D), volumetric (3D), and spatiotemporal (4D) measurements are outlined.

***Experimental design.**** Tissue handling, preservation, and spatial sampling introduce unique sources of bias and technical variation. This chapter discusses the tissue lifecycle, including fixation strategies, histological artifacts, FFPE-specific considerations, and statistical concepts such units of analysis and (pseudo-)replication. Additional topics include tissue microarrays (TMAs), targeted gene panel design, and region of interest (ROI) selection.

***Infrastructure.**** In Bioconductor, the primary class for handling **single-cell** data is *SingleCell-Experiment* [20] (SCE), which extends *SummarizedExperiment* by a series of characteristics specific to these data. *SpatialExperiment* [18] (SPE) extends SCE with additional customizations to store **spatial** information; this class is used in OSTA predominantly. SPE has been extended through *SpatialFeatureExperiment* [21] to further accommodate graphs and geometries. *MoleculeExperiment* [19] is an extension to SPE designed for iST in particular.

There are several **other frameworks** outside Bioconductor that (mostly) rely on in-house classes: *Seurat* [115] (R) and *Scanpy* [116] (Python) for single-cell; *Giotto* [117, 2] and *VoltRon* [23] (R), as well as *Squidpy* [3] and *SpatialData* [118] (Python) for spatial omics. Open-source **interactive visualization** tools include *Napari* [90] and, at the single-cell level, R/Bioc's *iSEE* [22]. Lastly, **commercial solutions** from Bruker, 10x Genomics, and Vizgen are also available (with the corresponding instruments).

***Ecosystem.*** Bioconductor hosts more than two thousand software and data packages for a variety of tasks and technologies. The *BiocPkgTools* [6] package allows for programmatic exploration of the ecosystem (e.g., package metadata, download statistics). Using *BiocViews* [5], we explore the number, lifetimes, and tasks of single-cell and spatial transcriptomics packages available.

***Importing.**** Running 10x Genomics' SpaceRanger on **Visium, Visium HD, and Xenium** data generates a set of standardized output files. For Visium (HD), these comprise raw measurements analogous to scRNA-seq, but at the spot-/bin-level and including spatial coordinates. Readers from several R/Bioc packages can be used to import data from raw files into R, including *VisiumIO*, *XeniumIO*, *SpatialExperimentIO*, and *SpatialFeatureExperiment*.

Bruker's AtoMx allows exporting different types of objects from **CosMx** data, including "flat" (human-readable) files that represent processed outputs (e.g., segmentation-derived counts). Both *SpatialExperimentIO* and *SpatialFeatureExperiment* support reading CosMx data into R.

***Datasets**** used throughout the book have been deposited through Bioconductor's *ExperimentHub* (EH), or an Open Science Framework (OSF) repository. These include ST datasets from representative commercial platforms across different tissue types from mouse and human; multi-sample analysis chapters also use imaging mass cytometry data.

OSF datasets may be queried and downloaded programmatically using the *OSTA.data* R/Bioc package (see also Table S1); otherwise, EH commands can be used. All data are managed on-disk using *BiocFileCache* to prevent costly re-retrieval of datasets across sessions.

***Interoperability.*** Key software enabling interoperability between R and Python include *reticulate* [30], which provides an R interface to Python, including support to translate between objects from both languages; and, *basilisk* [27], which facilitates Python environment management within the Bioconductor ecosystem. In addition, the Quarto publishing system can generate polyglot dynamic reports (OSTA is written using Quarto).

For single-cell/spatial omics in particular, *zellkonverter* [29] and *anndataR* [28] support conversion between Python's *AnnData* and R/Bioc's SCE, as well as reading and writing of .h5ad files. Notably, *anndataR* provides an R-native (R6) representation of *AnnData*, allowing for efficient and robust handling. *Rarr* [25] and *arrow* interface with .zarr and .parquet files, respectively, which are often used to store different ST data components; and, *alabaster* [26] can read and write language-agnostic file artifacts (e.g., for on-disk representation of SCE and *AnnData* objects).

#### II Sequencing-based platforms

***Reads to counts.**** sST platforms employ next-generation sequencing (NGS) to quantify gene expression. Spatial information is encoded during platform manufacture through barcodes, each associated with transcripts measured during sequencing; this association is present within the structure of the reads.

OSTA provides theory on **sequencing** in transcriptomics, including read structure and file formats, and outlines **processing** undertaken to convert a series of reads into a count matrix. Complementing commercial software, the R/Bioc package *Rsubread* [36] serves read alignment and quantification; and, *stPipe* [35] is a platform-agnostic tool for unified processing of sST data.

***Quality control*** (QC) aims to remove low-quality observations or technical artifacts to mitigate noise and bias in downstream analyses. Based on a set of observation-level QC metrics (e.g., number of uniquely detected features), **global outliers** are typically identified by applying thresholds to sample-level QC metrics (e.g., using *scater* [37]).

This approach is standard for scRNA-seq data, but assumes that metrics are independent of biology, which is commonly not adequate in ST [119]. Instead, *SpotSweeper* [39] identifies **local outliers** by comparing a given observation's metrics to those of its local neighborhood.

***Deconvolution.*** sST data can contain zero to multiple cells per observation, which might cover them fully or only partially. As a result, per-spot/bin observations can represent a mixture of transcriptional programs; deconvolution aims at estimating the contribution of different cell types to each observation.

Dozens of methods have been proposed for this purpose. In OSTA, we showcase the use of R/Bioc packages *CARDspa* [74] and RCTD [73] (*spacexr* package); the latter has been shown to perform particularly well in independent benchmark studies [120, 121, 122]. Notably, deconvolution has also been applied to provide an estimate of spatial 'bleeding' in iST data [69].

***Workflows.*** The first workflow chapter analyses postmortem human brain tissue from the dorsolateral prefrontal cortex (DLPFC) region [8]. We also showcase the complementary *SpatialLIBD* [33] R/Bioc package, which provides interactive analysis capabilities using *Shiny*.

Remaining workflows use data on a human colorectal cancer (CRC) biopsy from [7] (different sections subjected to Visium and Visium HD). The Visium workflow (Figure S4) recapitulates QC, unsupervised clustering and deconvolution, quantification of hallmark gene sets, and various exploratory analyses. For Visium HD, we perform bin- (Figure S5) and cell-level (Figure S6) analysis, the latter using H&E-segmented data bundled with standard SpaceRanger v4 outputs.

#### III Imaging-based platforms

***Segmentation.**** iST relies on microscopy stains to estimate (nucleus or membrane) boundaries, and assign molecular readouts to their cell of origin. Such readouts can be discrete (e.g., molecules) or continuous (e.g., fluorescence). The resulting measurement matrix of features (e.g., genes or proteins) $\times$ cells forms the basis for numerous analysis tasks.

OSTA briefly summarizes segmentation **approaches** (image-based [67], transcript-based [66], hybrid [65]). In addition, the phenomenon of spatial **bleeding** is discussed [72], including methods for decontamination (e.g., *FastReseg* [71] in R, *segger* [70] in Python). Lastly, **segmentation-free** analyses are noted (e.g., *SSAM* [68]).

***Quality control.*** Standard QC metrics (see above) can also apply to data from iST, however, a few considerations should be made: iST are often targeted (an incomplete representation of a cell's transcriptome); ribosomal or mitochondrial targets are typically lacking; and, data are acquired through iterative imaging of predefined regions, so-called fields of view (FOVs).

Then again, iST platforms include morphology information (e.g., cell area and eccentricity) and information from IF stains (e.g., of nuclei and membrane). The R/Bioc package *SpaceTrooper* [40] combines various QC metrics into a quality score, incorporating iST-specific metrics in particular; aberrant cells may be excluded, or flagged for extra considering downstream.

***Neighborhood analysis*** aims to investigate the composition of and interaction between proximal cells and cell types. Many spatial analyses rely on identifying **nearest neighbor** (NN) cells. OSTA recommends *RANN* for this purpose, which finds NNs in $O(n \log n)$ time for $n$ cells (conventional approaches would take $O(n^2)$ time) via a C++ backend, and supports fixed-radius searches.

Spatial **niche analysis** aims to identify regions of homogeneous microenvironments. The R/Bioc package *imcRtools* [53], for instance, clusters cells based on cell type frequencies in Euclidean neighborhoods. Complementarily, **co-localization** may be investigated using *hoodscanR* [52] to quantify attraction/repulsion between cell types as well as local mixing (e.g., entropy).

Spatial **density analysis**, as provided by *scider* [51], uses Kernel density estimation to describe the spatial distributions of cells and define regions of interest (ROIs); tertiary lymphoid structures, for example, correspond to aggregates of specific immune cell types. Density estimates can be used to evaluate co-localization globally; to quantify trends in tissue organization (e.g., remission of immune cells with increasing tumor density); and, ROIs may be subject to a number of downstream analyses, including DE testing (e.g., with *limma* [123], *edgeR* [124], and *DESeq2* [125]).

***Cell-cell communication*** (CCC). Besides spatial patterns in cellular organization and gene expression, it is further of interest to characterize how cells coordinate activity, e.g., through protein-mediated signaling.

In OSTA, we showcase the use of Python's *COMMOT* [126], an optimal transport-based and spatially aware tool for CCC inference that can link to databases of signaling interactions and pathways (e.g., *CellPhoneDB* [127]). This is done using *reticulate* and *zellkonverter* for seamless integration (setup in R, running *COMMOT* in Python, and exploring results back in R).

***Sub-cellular analysis**** aims to identify and quantify intra- and/or extra-cellular patterns and compartmentalization of transcripts (e.g., nucleus and cytoplasm of a cell). OSTA gives an overview of methods in this realm, including *SpatialFeatures* in R, and *Bento* [128], *CellSP* [129], *ClusterMap* [130], *FISHFactor* [131] *SpaGNN* [132] in Python.

***Workflows.*** An exemplary Xenium workflow (Figure S7) uses the same CRC dataset as for sST workflows (see above). The CosMx workflow (Figure S8) relies on 1K-plex data on mouse brain tissue, and includes data type-specific QC, label transfer-based annotation, as well as exemplary downstream analyses (namely, identification of SVGs based on spatial neighborhoods, and spatial statistics to investigate cell-cell interactions).

#### IV Platform-independent analyses

***Normalization*** aims to remove technical variability while preserving biological signals. In scRNA-seq, scaling-normalization aims at removing differences in sequencing coverage between libraries (*scater* [37] and *scran* [78] provide global and cell type-specific approaches, respectively). For ST data, however, total counts have been observed to be confounded with biology [133, 119].

The R/Bioc package *SpaNorm* [38] incorporates spatial information alongside gene expression to decompose variation into a technical and biological component. Because iST are most often targeted (and sequencing-free), area-normalization might be more appropriate still. In OSTA, we explore scaling-, area-, and spatially aware normalization, including qualitative comparisons.

***Dimensionality reduction, clustering & annotation.*** In single-cell omics data analysis, dimensionality reduction (DR) techniques are often categorized as linear (e.g., PCA), or non-linear (e.g., t-SNE, UMAP); *scater* [37] implements these and additional methods. For clustering, *BiocNeighbors* and *scran* [78] implement various algorithms for graph construction (e.g., $k$NN) and community detection (e.g., Leiden [134]), respectively.

In the context of ST data, we instead distinguish between spatially aware and non-spatial methods, i.e., whether or not DR/clustering incorporates physical locations. Throughout OSTA, we demonstrate both spatial DR combined with non-spatial clustering (using *BANKSY* [41]), and non-spatial DR combined with spatial clustering (using PCA and *BayesSpace* [42]). In addition, label transfer approaches that rely on corresponding single-cell reference data are demonstrated (using *SingleR* [44]). In conclusion, the chapter discusses strategies for annotation (from manual to (semi-)supervised to foundation models), reference resources (e.g., consortia efforts), and how to assess both (i.e., reference suitability and prediction quality).

***Feature selection & testing.*** OSTA distinguishes three broad types of features, namely: highly variable genes (HVGs), spatially variable genes (SVGs), and differentially expressed genes (DEGs). Though defined differently, these are highly connected from a biological perspective and thus discussed jointly.

For example, *scran* [78] models the mean-variance relationship across all observations, decomposing variance into a technical and biological component; **HVGs** are selected based on the latter, and commonly serve as input for DR and clustering. **DEGs** can be identified through pair-wise statistical tests between clusters to find markers (*scran* offers various options), or by testing for state changes across conditions (*muscat* [135], *miloDE* [136], and *lemur* [137] implement different approaches). Numerous tools to identify **SVGs** – each different in terms of methodology – have been proposed (e.g., R/Bioc packages *nnSVG* [79] and *DESpace* [80]) and benchmarked [138, 139].

***Feature-set signatures.*** Instead of identifying relevant features *de novo*, we can evaluate gene sets known to orchestrate biological functions or pathways (e.g., metabolism, cell cycle and death). Approaches to do this rely on **databases** of transcription factor binding sites, gene regulatory networks, or annotated gene sets (e.g., MSigDB [140], to which the R/Bioc package *msigdb* provides a programmatic interface); such sets may also stem from the literature or in-house studies.

R/Bioc packages for set **scoring** include *AUCell* [77] and *UCell* [141], which both rely on a rank-based approach to quantify the enrichment of query genes. Besides cell-/bulk-level quantification, a variety of **downstream** analyses can bring additional biological insights: scoring results may be summarized at different levels (e.g., clusters, niches, samples); they may be correlated with each other or orthogonal readouts (e.g., from signaling or trajectory inference); and, they may be compared across experimental conditions.

***Spatial statistics.*** Historically, the R programming language has been dedicated to statistical computing. Its application to spatial data dates back decades, primarily in epidemiological and geospatial research. As a result, various tools for spatial analyses have been established, including *spatstat* [84], *sp* [83], and *sf* [82].

OSTA recapitulates key concepts from *pasta* [85]: an extensive resource that provides theoretical background and practical examples for incorporating spatial statistics in spatial omics data analysis. *pasta* distinguishes two streams of data and corresponding types of analyses, namely, **point process** and **lattice** data. Analyses include but are not limited to: spatial autocorrelation, gene co-expression, attraction/avoidance of cells, cellular co-localization, etc.

***Image analysis.**** ST data can include biological images, such as immunofluorescence (IF) and hematoxylin and eosin (H&E) stainings. Ultimately, we aim to extract quantitative features from such images, and to incorporate them with molecular readouts.

Open-source software for visualization, annotation, and quantitative analysis of H&E and IF images include QuPath [91] and *Napari* [90]. Important resources include extensive collections of biomedical cancer images (e.g., TCGA and TCIA), and the histopathology foundation model Prov-GigaPath [92]. *scikit-image* and *Squidpy* [3] implement methods to extract quantitative image features in Python. Lastly, the R/Bioc package *imageTCGA* provides a programmatic interface for exploring TCGA-derived data.

***Deep learning.**** Deep learning has become an increasingly important component of spatial omics, supporting applications ranging from image analysis and cell segmentation to spatial domain identification and cell type annotation.

OSTA introduces the principles of modern deep learning, including convolutional, graph, and transformer neural networks, before motivating the emerging paradigm of foundation models. The chapter surveys representative foundation models for histology (H&E), multiplexed immunofluorescence (IF), single-cell, and ST data, discusses strategies for adapting pre-trained models (zero-shot inference, linear probing, and fine-tuning), and outlines practical considerations (e.g., reproducibility and computational requirements).

#### V Multi-sample analyses

***Differential spatial patterns.*** When multi-sample, multi-condition ST data are available, differential analysis can be performed to identify genes with spatial expression patterns that change between conditions; OSTA demonstrates the use of *DESpace* [80] to identify such genes.

***Differential co-localization.*** Similarly, one can investigate how co-localization of cell types changes across experimental conditions; here, the R/Bioc packages *spicyR* [87] and *spatialFDA* are showcased, which rely on spatial statistic curves (e.g., Ripley's $K$ function) to model co-localization patterns, and linear (mixed) models and functional data analysis for inference, respectively.

***Structure-based analysis*** focuses on reconstructing and quantifying anatomical structure-derived features (e.g., pancreatic islets). The R/Bioc package *sosta* [86] provides various methods for point pattern-based segmentation of such structures, and metrics for their quantification (e.g., shape statistics). In multi-sample settings, structures may be subject to comparison across experimental conditions. Because they represent repeated measurements, linear mixed models can be used to account for correlations (e.g., using *lme4* [142] and *lmerTest* [143]).

#### VI Cross-platform analyses

***Registration*** is defined as the task of mapping features or spatial locations of observations from a query to a reference assay. In the context of ST, we distinguish two types of registration. Spatial **alignment** is typically image-based; OSTA provides an example of using *VoltRon* [23] to align Xenium data with a post-run H&E staining. Omics-based approaches that depend only on the spatial distribution of molecule profiles are summarized as well (e.g., *PASTE* [95] and *SLAT* [94] in Python). Spatial **reconstruction** leverages spatially-resolved reference data to reconstruct the spatial coordinates of query single cells; methods mentioned include *CeLEry* [96], *novoSpaRc* [99], and *SpaOTsc* [64] in Python.

***Imputation.*** Most commonly employed iST assays resolve a limited number of features. Meanwhile, non-spatially resolved assays are of whole-transcriptome plexity. One can integrate spatially resolved data with scRNA-seq data using common features, and predict missing features using transcriptionally similar cells.

OSTA showcases using *harmony* [101] for integrating low-plex iST with scRNA-seq data in order to impute transcriptome-wide measurements. Other methods for this task include *LIGER* [93] in R and *Tangram* [98] in Python.

***Workflow.*** An exemplary workflow chapter uses Visium and Xenium data on human breast cancer from [9]. These are used to showcase how to spatially align, integrate (using *harmony* [101]), and impute data across modalities (via smoothing across cells that are transcriptionally similar based on shared features).

#### VII Beyond this book

OSTA concludes with a part that consolidates analytical concepts and tasks not explicitly covered in the current version of the book. At the time of writing, these include detailed methodological foundations (e.g., mathematics), batch correction and data integration, trajectory inference, copy number variation (CNV) inference, and multi-modal data analysis.

Each section provides a brief overview of the topic, together with curated pointers to relevant literature and resources. This part therefore acknowledges current limitations while serving as a foundation for future contributions from community members with expertise in these areas.

48. Domínguez Conde C, Xu C, Jarvis LB, Rainbow DB, Wells SB, Gomes T, Howlett SK, Suchanek O, Polanski K, King HW, Mamanova L, Huang N, Szabo PA, Richardson L, Bolt L, Fasouli ES, Mahbubani KT, Prete M, Tuck L, Richoz N, Tuong ZK, Campos L, Mousa HS, Needham EJ, Pritchard S, Li T, Elmentaite R, Park J, Rahmani E, Chen D, Menon DK, Bayraktar OA, James LK, Meyer KB, Yosef N, Clatworthy MR, Sims PA, Farber DL, Saeb-Parsy K, Jones JL, and Teichmann SA. Cross-tissue immune cell analysis reveals tissue-specific features in humans. Science (New York, N.Y.) 2022; 376:eabl5197. DOI: 10.1126/science.abl5197

83. Pebesma E and Bivand RS. Classes and Methods for Spatial Data: the sp Package. The Newsletter of the R Project 2005; 5

90. Sofroniew N, Lambert T, Bokota G, Nunez-Iglesias J, Sobolewski P, Sweet A, Gaifas L, Evans K, Burt A, Doncila Pop D, Yamauchi K, Weber Mendonça M, Liu L, Buckley G, Vierdag WM, Monko T, Willing C, Royer L, Can Solak A, Harrington KIS, Abramo J, Ahlers J, Althviz Moré D, Amsalem O, Anderson A, Annex A, Aronssohn C, Balzaretti F, Boone P, Bragantini J, Bussonnier M, Caporal C, Coccimiglio I, Eglinger J, Eisenbarth A, Freeman J, Gohlke C, Gunalan K, Halchenko YO, Har-Gil H, Harfouche M, Hilsenstein V, Hutchings K, Kozar R, Lauer J, Lichtner G, Liu H, Liu Z, Lowe A, Marconato L, Martin S, McGovern A, Migas L, Miller N, Miñano S, Muñoz H, Müller JH, Nauroth-Kreß C, Obenhaus HA, Palecek D, Pape C, Perlman E, Pevey K, Peña-Castellanos G, Pierré A, Pinto D, Rodríguez-Guerra J, Ross D, Russell CT, Ryan J, Selzer G, Smith M, Smith P, Sofiiuk K, Soltwedel J, Stansby D, Vanaret J, Wadhwa P, Weigert M, Windhager J, Winston P, and Zhao R. napari: a multi-dimensional image viewer for Python. Zenodo 2025. DOI: 10.5281/zenodo.16883660
